# Deep learning detection of dynamic exocytosis events in fluorescence TIRF microscopy

**DOI:** 10.1101/2024.09.09.611975

**Authors:** Hugo Lachuer, Emmanuel Moebel, Anne-Sophie Macé, Arthur Masson, Kristine Schauer, Charles Kervrann

**Affiliations:** Université de Paris, CNRS, Institut Jacques Monod, 75013 Paris, France; SAIRPICO team, Centre Inria de l’Université de Rennes, INSERM-U1143, Institut Curie, PSL Research University, Campus Universitaire de Beaulieu, Rennes Cedex, France; Cell and Tissue Imaging Facility (PICT-IBiSA), Institut Curie, PSL Research University, Paris, France; Tumor Cell Dynamics Unit, Inserm U1279 Gustave Roussy Institute, Université Paris-Saclay, Villejuif 94800, France

## Abstract

Segmentation and detection of biological objects in fluorescence microscopy is of paramount importance in cell imaging. Deep learning approaches have recently shown promise to advance, automatize and accelerate analysis. However, most of the interest has been given to the segmentation of static objects of 2D/3D images whereas the segmentation of dynamic processes obtained from time-lapse acquisitions has been less explored. Here we adapted DeepFinder, a U-net originally designed for 3D noisy cryo-electron tomography (cryo-ET) data, for the detection of rare dynamic exocytosis events (termed ExoDeepFinder) observed in temporal series of 2D Total Internal Reflection Fluorescent Microscopy (TIRFM) images. ExoDeepFinder achieved good absolute performances with a relatively small training dataset of 60 cells/∼12000 events. We rigorously compared deep learning performances with unsupervised conventional methods from the literature. ExoDeepFinder outcompeted the tested methods, but also exhibited a greater plasticity to the experimental conditions when tested under drug treatments and after changes in cell line or imaged reporter. This robustness to unseen experimental conditions did not require re-training demonstrating generalization capability of ExoDeepFinder. ExoDeepFinder, as well as the annotated training datasets, were made transparent and available through an open-source software as well as a Napari plugin and can directly be applied to custom user data. The apparent plasticity and performances of ExoDeepFinder to detect dynamic events open new opportunities for future deep-learning guided analysis of dynamic processes in live-cell imaging.

## Main

Technological improvements in imaging accelerate the amount of acquired data. This situation requires new methods to automatically extract the tremendous quantity of information present in them. It is clearly established that Deep-Learning-based image segmentation methods, and especially U-net^1^, surpass conventional techniques and exhibit a remarkable generalisation capacity^1,2^. However, most of the studies are restricted to the segmentation of static biological objects. Deep-Learning-methods are rarely applied to dynamic processes^3^ while several model-based methods have been developed in the past decades^4–7^. Here we focus on the supervised Deep-Learning-assisted detection of rare dynamic exocytosis events (< 1 event per frame in average) in videos. Exocytosis is the fusion of an intracellular vesicle with the plasma membrane to release its content. Many conventional methods of exocytosis detection have been already published^8–23^ including a deep-learning strategy^24^. These methods detect exocytosis events visualized by an exocytic reporter protein tagged with a pH-sensitive fluorophore^25^ imaged in Total Internal Reflection Fluorescence Microscopy (TIRFM). In general, a sudden peak of fluorescence intensity (termed “puff”) followed by an exponential decay of the signal (**F1A**) is detected. Unfortunately, these methods are often poorly evaluated on benchmark datasets and are not publically available.

Here we present an adaptation of DeepFinder^26^ (**see Methods**), a U-net deep-learning method, for the detection of exocytosis events in large 2D+Time volumes. DeepFinder was originally developed for the identification of macromolecules in 3D noisy cellular cryo-tomograms (cryo-ET) and is considered as a top-rank method, confirmed in several SHREC challenges^27^. To train ”ExoDeepFinder” we took advantage of our recently published large dataset^28^ monitoring lysosomal exocytosis events via imaging of VAMP7-pHluorin by TIRFM. The coordinates of all exocytosis events were manually annotated by a single expert based on the characteristic “puff” signature (**F1A-B and movie S1**). Because DeepFinder performances for cryo-ET segmentation substantially increased thanks to a multi-class strategy, we applied a similar approach here: We combined manual annotations of exocytosis events with automatic algorithm-generated annotations of docked vesicles at the plasma membrane^29^ that form bright foci without fusion with the plasma membrane (**see Methods**) (**Fig 1C**).

**Figure 1.**
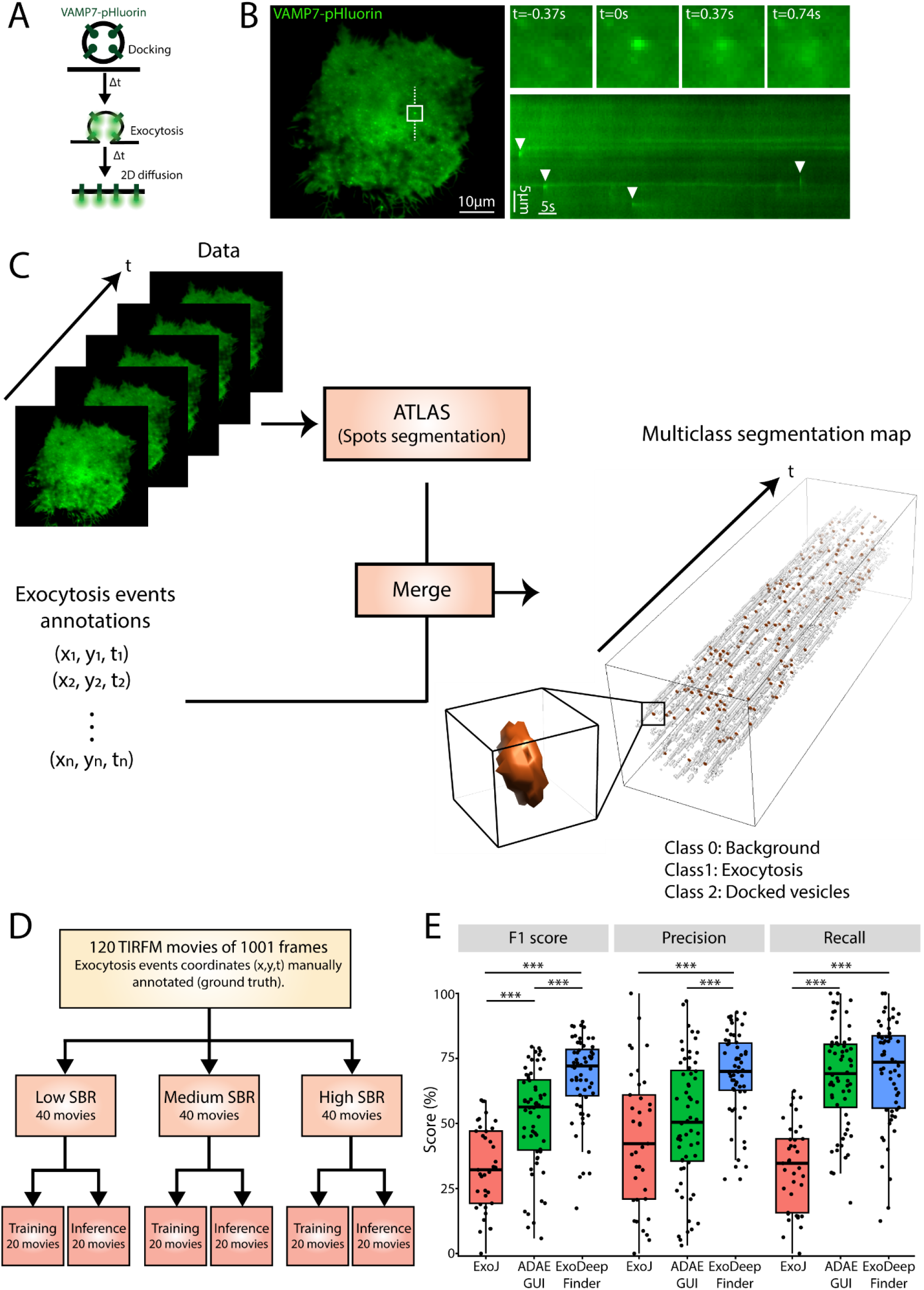
**A**. Schematic representation of the exocytosis of a VAMP7-pHluorin-positive vesicle: the low pH of the acidic lumen quenches the fluorescence of pHluorin. During exocytosis, protons are released and pHluorin starts to emit light. An exocytosis event is followed by the 2D diffusion of VAMP7-pHluorin at the plasma membrane. **B**. TIRFM image of VAMP7-pHluorin in a transfected RPE1 cell. The inset represents the field in the white square showing one exocytosis event at different time points, *t*=0 represents the peak of the exocytosis event. A kymograph is plotted along the dashed white line and arrowheads indicate several observed exocytosis events. **C**. Workflow of the data preparation for ExoDeepFinder training. On the one hand, manual annotation gives the coordinates of ‘ground truth’ exocytosis events, on the other hand, spots corresponding to docked vesicle are detected thanks to the ATLAS algorithm. Both annotations are merged to produce a multiclass segmentation map. The resulting spatial coordinates are converted into a 3D mask. The luminescence of an exocytosis event is isotropic in the (*x,y*) plane and has an exponential decay in t. Therefore, we model an exocytosis event as a tube with an exponentially decaying radius starting at R=4 pixels and ending at R=1 pixel, the length of the tube being 3 frames in the temporal dimension as illustrated in the insight. Hence, our segmentation map is composed of 3 classes: background (class 0), *bona fide* exocytosis event (class 1) and docked vesicles/constant spot (class 2). **D**. Organization of the dataset. The dataset is composed of 120 TIRFM movies of 1001 frames. Each movie has a manual annotation of exocytosis event coordinates constituting the ground truth. The dataset is split into 3 equal subgroups according to the average SBR (=F/F_0_) of each movie. Then, each SBR group is randomly split into two equal sub-groups, one dedicated to ExoDeepFinder training and the other one to inference *i*.*e*. ExoDeepFinder evaluation. **E**. Comparison of ExoDeepFinder performances with ExoJ and ADAE GUI on the same inference dataset of 60 movies. Significance has been evaluated with a Kruskal-Wallis test (p<0.001) and pairwise comparison with a Dunn’s post-hoc test with a multiple comparison Holm correction, ***p<0.001 (other comparison are not significant *i*.*e*. p>0.05). ExoJ has less number of movies analyzed (hence less total number of events) because the analysis cannot be performed for 30-40% of the data (**see method**). ExoDeepFinder was trained on the total 60 movies of the training dataset (model all).

**Figure 2.**
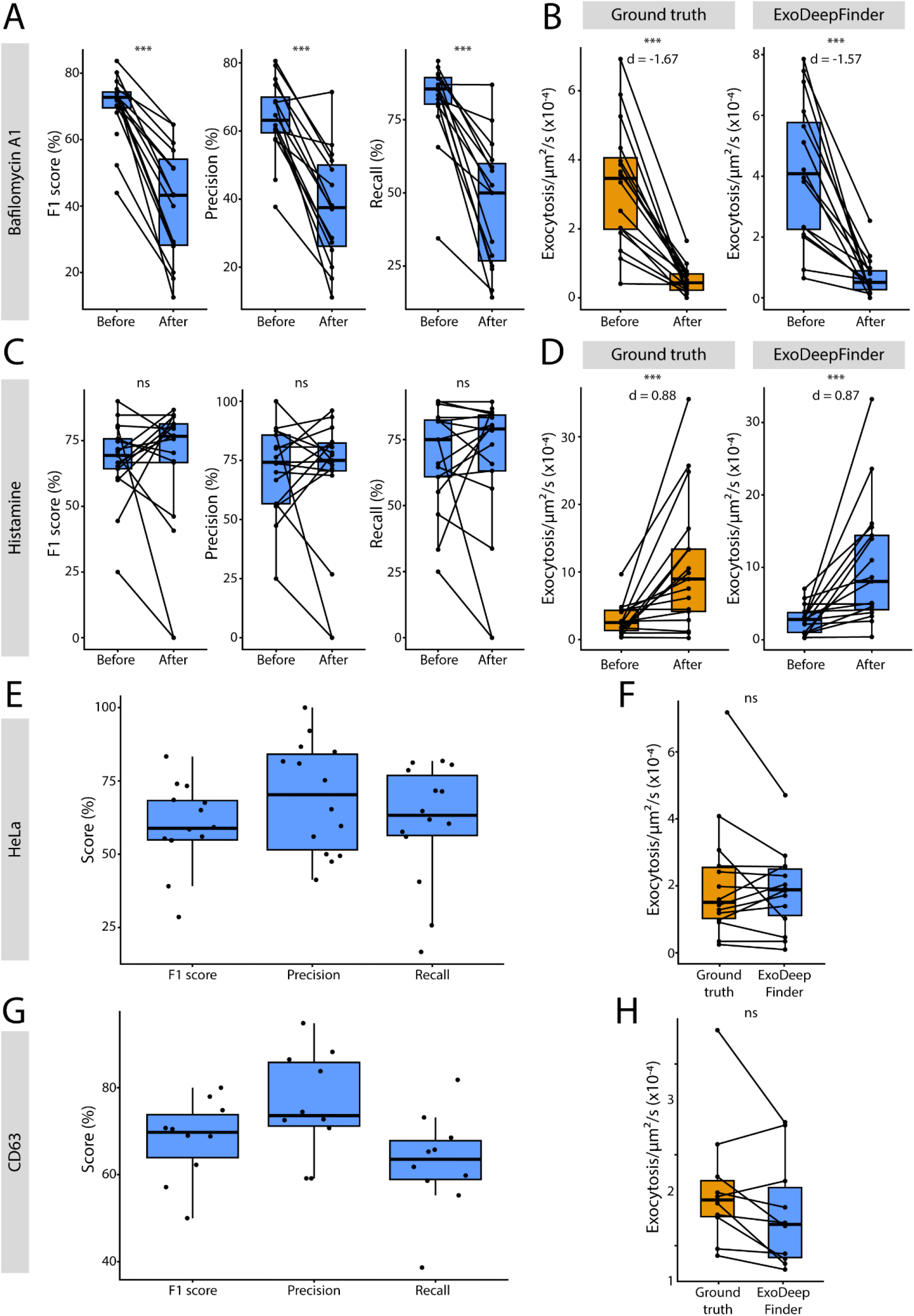
**A**. ExoDeepFinder performances before and after Bafilomycin A1 (100 nM, 60 min) treatment. Note that one point is unpaired that represents a cell with no predicted events after treatment (hence, F1-score, Precision and Recall cannot be defined). **B**. Exocytosis rate before and after Bafilomycin A1 (100 nM, 60 min) treatment measured by manual detection (ground truth) and compared to ExoDeepFinder detection. In A-B, n=16 cells from three independent experiments with a total 3008 and 401 exocytosis events before and after drug addition, respectively, are shown. **C**. ExoDeepFinder performances before and after histamine (100μM, cells immediately imaged) treatment. **D**. Exocytosis rate before and after histamine (10 μM, cells immediately imaged) treatment measured by manual detection (ground truth) and compared to ExoDeepFinder detection. In C-D, n = 17 cells from three independent experiments with a total 2292 and 3201 exocytosis events before and after drug addition respectively, are shown. **E**. ExoDeepFinder performances in VAMP7-pHluorin transfected HeLa cells. **F**. Comparison of the exocytosis rate measured by manual detection (ground truth) and compared to ExoDeepFinder detection. In E and F, 14 cells analyzed from a single experiment with a total of 1045 exocytosis events, are shown. **G**. ExoDeepFinder performances in CD63-pHluorin transfected RPE1 cells. **H**. Comparison of the exocytosis rate measured by manual detection (ground truth) and compared to ExoDeepFinder detection. In G and H, 10 cells analyzed from a single experiment with a total of 972 exocytosis events, are shown. In A-D, F and H significance has been evaluated with paired Wilcoxon test, ns p>0.05 and ***p<0.001. In B and D, effect sizes are measured with the Cohen’s *d* for paired samples. In A-H, ExoDeepFinder was trained on the total 60 movies of the training dataset (model all).

Furthermore, we evaluated detection performance as a function of the “Signal-to-Background Ratio” (SBR). We defined the SBR as a ratio between the local fluorescence intensity F after vesicle fusion with the plasma membrane and the fluorescence intensity F_0_ before this peak, *i*.*e. SBR* = *F*/*F*_0_. We constituted a dataset of 120 TIRFM movies (for a total of 20 567 exocytosis events) of 1001 frames (about 6 minutes) that we divided into three groups based on their average SBR (**F1D**). Then we split randomly each SBR group into two sub-groups, one dedicated to training of the network and the other one to its evaluation (**F1D**). We made our dataset entirely available (**see supplementary**).

We trained ExoDeepFinder on the three sub-datasets (low, medium, and high SBR values), on the combination of these three datasets (Subsets A to D) (**Table S1**) and on the full training dataset (i.e. 60 movies). ExoDeepFinder performances for each training set were evaluated in terms of F1-score, Recall and Precision over the whole inference dataset (*i*.*e*. 60 movies). As commonly observed for neural network approaches, the best performances were achieved with the training over the full training dataset with 67.64% for F1, 70.07% for Recall and 68.75% for Precision (**Table 1**). Moreover, we compared these performances to the two unsupervised, conventional exocytosis detection methods that are publically available i) ExoJ^22^ and ii) ADAE GUI^20,21^. We compared the method performances over 60 movies dedicated to inference (**F1D and movie S2**). ExoDeepFinder performed better than ExoJ and ADAE GUI in terms of F1-score and Precision. For Recall, ExoDeepFinder was statistically indistinguishable from ADAE GUI, but better than ExoJ (**F1E**). Note that despite numerous efforts to run the analysis on different computers overnight and with sub-parts of the movies, ExoJ failed to produce any results for a high percentage (30-40%) of the data (**S1E**) (**see Methods**). We noticed that ExoDeepFinder performances were robust over different frame rates (**S2B**) and correlated with the F/F_0_ SBR (**S2B**).

**Table 1.** Performances of ExoDeepFinder for the different training datasets defined in **Table S1**. Values are mean ± SD.

|  | F1-score (%) | Recall (%) | Precision (%) |
| --- | --- | --- | --- |
| All | 67.64 ± 15.91 | 70.07 ± 19.81 | 68.75 ± 16.57 |
| Subset A | 54.77 ± 20.50 | 52.30 ± 23.02 | 64.65 ± 22.13 |
| Subset B | 61.68 ± 20.06 | 62.75 ± 23.65 | 65.94 ± 20.35 |
| Subset C | 63.62 ± 19.82 | 63.95 ± 23.68 | 69.60 ± 20.30 |
| Subset D | 62.67 ± 19.39 | 62.51 ± 23.58 | 69.67 ± 20.23 |
| Subset E | 64.51 ± 19.93 | 62.95 ± 22.96 | 70.96 ± 19.51 |
| Low SBR | 62.19 ± 18.83 | 69.91 ± 21.23 | 61.54 ± 22.20 |
| Medium SBR | 58.26 ± 20.22 | 70.05 ± 23.07 | 58.31 ± 24.74 |
| High SBR | 60.37 ± 21.18 | 58.26 ± 23.16 | 68.67 ± 22.70 |

Next, we assessed ExoDeepFinder’s efficiency on cells treated with different drugs that interfere with secretion without retraining. We analyzed cells before and after treatment with Bafilomycin A1, a drug impairing lysosomal pH homeostasis that leads to decreased exocytosis rate and additionally changes cell morphology by creating stable and bright foci at the plasma membrane. ExoDeepFinder performances decreased significantly after the drug treatment when compared to same cells before treatment (**F2A**). However, ExoDeepFinder was still able to estimate correctly exocytosis rate and produced a correct estimation of drug effect size (**F2B**) even though it was exclusively trained on constitutive exocytosis events. Indeed Cohen’s *d* predicted by ExoDeepFinder was -1.57 while ground truth was *d*=-1.67. Similarly, ExoJ and ADAE GUI performances decreased (**S3A and C**), but contrarily to ExoDeepFinder, ADAE GUI and ExoJ made wrong estimations of exocytosis rates, especially after treatment (**S3B and D**). Both predicted an effect in the wrong direction (*d*=0.10 for ExoJ and *d*=0.37 for ADAE GUI).

Additionally, we analyzed cells treated with histamine that stimulates lysosomal exocytosis^28,30^. Contrarily to Bafilomcyine A1, cell morphology and the aspect of individual secretory events was preserved. In these conditions, we observed that ExoDeepFinder performances were not impaired (**F2C**) and estimated precisely exocytosis rate as well as effect size (predicted *d*=0.87 while ground truth *d*=0.88) (**F2D**). ExoJ and ADAE GUI performances were not impacted by histamine treatment either (**S3E and G**). However, due to their inherent lower performances, estimated exocytosis frequency and effect size were less accurate (*d*=0.37 for ExoJ and *d*=0.98 for ADAE GUI) (**S3F and H**).

We next detected VAMP7+ exocytosis events in another cell type (HeLa) still without re-training of ExoDeepFinder. The ExoDeepFinder performances were similar good as in RPE1 cells (**F2E**), and the estimated exocytosis rate was statistically indistinguishable from ground truth (**F2F**). Surprisingly, ExoJ performances decreased a lot (**S4A**): the estimated exocytosis rate decreased significantly compared to ground truth (**S4B**). This is surprising, because part of the ExoJ validation was performed on the VAMP7-pHluorin probe in HeLa cells^22^. ADAE GUI Precision decreased slightly (**S4C**), and the predicted exocytosis rate was significantly over-estimated (**S4D**).

Finally, we evaluated the detection of another cargo, CD63, a transmembrane protein of the tetraspanin family that is used as a marker of the secretion of multi-vesicular bodies. Exocytosis events in RPE1 cells were monitored with a CD63-pHluorin probe^30^. We found that ExoDeepFinder performances were roughly preserved (**F2G**), and exocytosis rate was statistically indistinguishable to ground truth (**F2H**). ExoJ F1-score and Recall substantially increased for this dataset but the Precision value decreased (**S4E**). Note that although CD63-pHluorin was used to validate ExoJ^22^, the exocytosis rate predicted by ExoJ was underestimated, even if not significantly (**S4F**) due to the low number of movies that were analyzable with ExoJ. Contrary, ADAE GUI F1-score and Precision decreased substantially (**S4G**), and the predicted rate was significantly over-estimated compared to ground truth (**S4H**).

In conclusion, we developed ExoDeepFinder using the U-net DeepFinder architecture, originally designed for macromolecule detection in 3D cryo-electron cell tomograms. We demonstrated that ExoDeepFinder, trained on 60 TIRF movies and hybrid annotations, outperformed unsupervised conventional methods of exocytosis detection (**F1E**) and was more robust (**F2**). Moreover, ExoDeepFinder achieved good performances even with smaller training datasets (**Table S1**). A multi-class training of ExoDeepFinder increased its performance^26^ (**F1C**), but may require additional work for the user such as the segmentation of counter-examples events. A particular advantage of ExoDeepFinder is its speed to analyze large 2D+time volume data: It takes about 30 seconds to process a video of 300 × 300 × 1000 voxels with no parameter adjustments, contrary to 10 to 20 minutes required with the two conventional image analysis algorithms used in our benchmark analysis that additionally required manual parameter calibration. The training required 8 to 18 hours of computing (once for all), depending on the desired number of epochs and the GPU performance. Importantly, we endeavored to produce an user-friendly software version of ExoDeepFinder publically avalable (**see data and software availability**). Moreover, our software allowed user to re-train the network on its own data. We showed that ExoDeepFinder is capable of imitating expert annotations on experimental videos. ExoDeepFinder was demonstrated to robustly perform on TIRF videos with variable experimental conditions (various cargo, cell lines, microscopes, cameras, fluorophores, SBRs, etc.) It is able to reliably detect exocytosis events on signals from previously unseen proteins. It can be retrained from an external dataset or fine-tuned to adapt to other target cell types or proteins. We provide an open-source implementation of the ExoDeepFinder software, the manual annotated dataset for training, and the trained model for inference and fine-tuning. ExoDeepFinder performances also open the possibility to develop similar Deep-Learning approaches dedicated to the analysis of other dynamic processes such as endocytosis or the detection of blinking spots in the context of super-resolution microscopy.

## Acknowledgments

We are grateful to Thierry Galli for the VAMP7-pHluorin plasmid and Jean Salamero for initial discussions on this project. We greatly acknowledge the Nikon Imaging Centre @ Institut Curie-CNRS, member of the French National Research Infrastructure France-BioImaging (ANR10-INSB-04). This work was supported by ARC (Association pour la Recherche sur le Cancer) PhD fellowship, FRM (Fondation Recherche Médicale) PhD extension fellowship, the ITMO Nanotumor grant and Myocortex grant from the Agence Nationale de la Recherche (ANR-21-CE13-0010-01) to KS and grants from the Labex Cell(n)Scale (11-LBX-0038), the Idex Paris Sciences et Lettres (ANR-10-IDEX-0001-02 PSL) and the National Research Agency (Increased ANR-20-ASTR-0005, France-BioImaging ANR-10-INBS-04-07).

