## Supplementary figures and text for "Deep learning detection of dynamic exocytosis events in fluorescence TIRF microscopy"

#### Supplementary text

##### Data and software availability

We publically shared our dataset (https://doi.org/10.5281/zenodo.11204932) including all the raw TIRFM movies in .TIF files as well as the coordinates of exocytosis events from manual annotation under .txt files representing ImageJ results (x,y,t coordinates are given by "X", "Y" and "Slice" columns).

The code of ExoDeepFinder can be downloaded for free from our GitHub website: https://github.com/deep-finder/tirfm-deepfinder along with accompanying documentation. Our Github hosts an user-friendly software version of ExoDeepFinder with a Graphical User Interface (GUI). Our Github also hosts a Python version of ExoDeepFinder than can be optionally run also with a GUI. ExoDeepFinder can be executed with scripts using the API (examples are provided) or with a GPU. To implement ExoDeepFinder, we used Keras (http://keras.io), an open-source toolbox written in Python and using the TensorFlow framework. All training procedures were achieved using a Nvidia A100 running CUDA 11.4.

##### Definition of the SBR and correlation with performances

Signal-to-Background Ratio (SBR) can be defined in two ways for our TIRFM movies, i) as a classic ratio between cell signal over background and ii) as a ratio between the local fluorescence intensity F after vesicle fusion with the plasma membrane and the fluorescence intensity $F_0$ before this peak (*i.e.* $SBR = F/F_0$). These two SBR are not correlated (**S1A**). We relied on the second SBR, because it directly characterizes single exocytosis events and is the usual feature detected by former algorithms. However, we noticed afterwards that ExoDeepFinder performances, similarly to ExoJ and ADAE GUI, correlated also with the first type of SBR (**S1B-D**). ExoJ performances correlated with both SBR as well as with the frame rate (**S1B**). Recall from ExoJ correlated positively with the frame rate (Spearman correlation coefficient ρ=0.22) but Precision correlates negatively with frame rate (Spearman correlation coefficient ρ=-0.36). ADAE GUI performances did not correlate with the frame rate but positively with both SBR, similarly to ExoDeepFinder (**S1C**).

##### Optimization of ExoJ

ExoJ detection method relies on a first step of spot detection by applying the "à trous" wavelet transform similarly to Yuan et al.[1]. Secondly, spots are tracked using the nearest-neighbour method, and finally, fusion events are predicted according to a peak of fluorescence followed by an exponential decay. ExoJ has free parameters (**see Methods**) that substantially impacted performance on our dataset: i) the wavelet scale σ played a role on the spot detection step, ii) the minimal $R^2$ tolerated fluorescence decay fitting and exocytosis spot radius fitting with a Gaussian and iv) an intensity ratio (dF/σ) was similar to $F/F_0$. We screened the ExoJ performances for the aforementioned parameters (**see Methods**). Surprisingly ExoJ failed to produce results for a high percentage (30-40%) of the data

(**S1E**) despite our effort to run the analysis on different computers overnight and with sub-parts of the movies. We identified the best set of parameters of ExoJ as those that maximize the average of the 3 scores: σ=5, decay $R^2$ = 0.8, radius $R^2$ = 0.7 and dF/σ =3.8 (**S1F**). This optimization of ExoJ performances allowed to perform a fair comparison with ExoDeepFinder.

#### Analysis of TP, FP and FN events

To characterize more precisely the differences in performances between methods, we examined the intersection of True Positive (TP), False Positive (FP) and False Negative (FN) events (**S2A**). Surprisingly, The count of TP events specific to ExoJ was extremely low (even taking into account that the total number of movies analyzed was lower for ExoJ) demonstrating that it only detected obvious events also recognized by ADAE GUI and ExoDeepFinder. ExoDeepFinder had the highest specific count of TP events highlighting that learning approaches are able to detect subtle and less stereotyped events. In terms of FP events, the intersections were quite small (and could potentially represent ambiguous events or even annotation errors) demonstrating that each method has a different way to "hallucinate" events. Still, the count of FP events specific to ExoJ was high suggesting that it missed many events and explained its low performances. Contrary, the count of FP events specific to ExoDeepFinder was low indicating that it mainly failed to detect events that were ambiguous or represented errors in annotation. We investigated how TP, FP and FN events differed in terms of SBR (defined as $F/F_0$) and exponential decay lifetime. We found that TP events were characterized by a higher SBR than FP and FP events for all 3 methods (**Table S2**). However, ExoDeepFinder showed the lowest average SBR among TP events suggesting that it additionally relied on other information. Similarly, long events were easier to detect, illustrated by the fact that TP events have longer fluorescence lifetimes for all 3 methods (**Table S3**). Interestingly, ExoDeepFinder and ADAE GUI sometimes miss-interpreted events with a long fluorescent exponential decay (*i.e.* high life-time) as exocytosis, contrarily to ExoJ which more often confused exocytosis with fast decay events (*i.e.* small life-time). Note that for both parameters, SBR and lifetime, ExoDeepFinder had a lower average value than ExoJ and ADAE GUI, suggesting a higher sensibility. Finally, we compared the methods in terms of localization precision of TP events (**see Methods**). ExoDeepFinder and ADAE GUI showed similar spatial localization precision, which was better than those of ExoJ (**S2B**). In terms of temporal Precision, ExoJ and ADAE GUI had similar performances, which yet were lower than that of ExoDeepFinder (**S2C**).

### Methods

#### Cell Culture

hTERT-immortalized retinal pigment epithelial cell line (hTERT RPE-1) were cultivated in DMEM/F12 media (Gibco, catalog # 21041-025) complemented with 10% Fetal Bovine Serum (Eurobio, catalog # CVFSVF00-01) (without antibiotics). HeLa immortalized cell line were cultivated in DMEM high glucose (4.5g/l) media (Gibco, catalog #11965092) (without antibiotics). Cells were maintained at 37°C with 5% $CO_2$ in a humidified incubator.

#### Transfection

Cells were transfected with the following constructs: VAMP7-pHluorin[2] and CD63-pHluorin (addgene plasmid #130901)[3]. Moreover our dataset includes VAMP7-pHluorin transfected cells co-transfected with mCh-Rab6A or Paxillin-mCh[4]. Cells were transfected with 800ng (or 2×400ng for co-transfection) of DNA using the JetPrime kit (Polyplus). Cells were imaged 24h after transfection.

#### Drug treatments

Cells were treated with Bafilomycin A1 (MedChemExpress, catalog # HY-100558, 100nM) for 1h before imaging or histamine (Merck, catalog # H7125, 100µM) and immediately imaged. For all drug conditions, a paired design was used: the same cell was imaged before and after the treatment. Note that for histamine treatment only the first 60s after drug addition were analyzed, because the effect of histamine on secretion rate was immediate but transient[5].

#### Micropatterning

Our dataset included some cells seeded on a micropatterned substrate. We followed the photolithography micropatterning protocol from Azioune et al.[6]. Briefly, coverslips (1.5H ThorLabs, Catalog # CG15XH1) were oxidized by plasma-cleaner (Harrick Plasma) during 5min. Coverslips were PEG-coated by incubating them on a drop of PLL-g-PEG [Surface Solutions, PLL(20)-g[3.5]- PEG(2)] (0.1 mg/mL diluted in water, 10mM HEPES, pH=7.4) in a moiety chamber during 1h. After coating, patterns were printed using a deep UV lamp (Jelight Company Inc, catalog # 342-220) with radiation passing through a photomask (DeltaMask) during 5min. Finally, patterns were fibronectin-coated by incubating coverslips on a drop of fibronectin (Merck/Sigma, catalog # F1141) (50 µg/mL diluted in water) and fibrinogen-Alexa647 (Molecular Probes, Invitrogen, catalog # F35200) (or fibrinogen-Alexa488) (5 µg/mL) in a moiety chamber during 1h. Coverslips were conserved at 4°C in PBS.

Cell seeding on micropatterns was described in Lachuer et al.[7]. Briefly, coverslips were maintained in magnetic chamlides for live imaging. ~200,000 trypsinized (Thermo Fisher, catalog # 12605010) cells were added in the chamlide chamber. After 10min incubation in 37°C incubator, cells were attached to the substrate. Cells were washed using DMEM/F12 media with 20mM HEPES (Gibco, catalog # 15630-056) with 2% penicillin/streptomycin (Gibco, catalog # 15140-122). Cells were incubated at least 3h in the incubator until full spreading on the micropattern. Cells were imaged the same day. Two different geometries of micropatterns were used: i) ring-shaped micropatterns with a diameter of 37µm and 7µm thickness of the adhesive ring; and ii) rectangular micropatterns with 9x40µm dimensions.

#### TIRFM

Non-patterned cells were seeded in fluorodishes (World Precision Instrument) coated with fibronectin (or Poly-L-Lysin (PLL) (Merck P4707) for some cells of the dataset). Media supplemented with 20mM HEPES was used for imaging. The acquisition was made using an inverted Nikon TIRFM equipped with an EMCCD camera (efficiency 95%) with a 100× objective (pixel size = 0.160 µm) with the 491nm laser. Time-lapse of VAMP7/CD63-pHluorin was acquired with a theoretical frame rate of one image every 300ms during 5min. Frame rate was set according to the half-life of exocytosis events. Due to microscopic device delay, the actual frame rate has been computed using the computer time of saved files.

#### Statistical Analysis

All statistical analyses were made with R [R Core Team (2021)]. The number of cells and the number of independent repetition is indicated in the legend. Since our cells were mostly isolated when imaged, we performed only single cells analysis, each cell was considered as independent, setting the sample size. The statistical test used was indicated in the legend. Tests were always conducted in a two-sided manner and a multiple comparison correction was applied when needed. To make no hypothesis about data normality we only applied nonparametric tests (Kruskal-Wallis test with Dunn's post-hoc test realised (with a Holm-Bonferroni correction) with the dunn.test package) and applied pairing when possible (paired Wilcoxon test). Finally, correlation was measured by Pearson correlation coefficient and tested with a t-test.

Effect size of paired data was analyzed with Cohen's *d*. Briefly, a value of exocytosis rate $x$ was obtained before and after treatment for each cell. The pairwise difference $\Delta x_i$ for cell $i$ was computed as $x_{after,i} - x_{before,i}$. The Cohen's *d* was computed as:

$$d = \frac{\overline{\Delta x}}{s_{\Delta x}}$$

With $\overline{\Delta x}$ the average of the pairwise differences $\Delta x_i$, and $s_{\Delta x}$ the standard-deviation of the pairwise differences $\Delta x_i$.

#### Image Analysis

##### ***Manual exocytosis detection (Ground truth)***

Exocytosis events were detected manually based on the visual recognition of a sudden burst of intensity signal followed by an exponential decay. However, this task is prone to error due to the size of the dataset and the ambiguity of some events. Our dataset probably includes FP and FN. However, these errors are minor, reflect real situations and did not impair ExoDeepFinder training.

##### ***Exocytosis detection with ExoJ***

ExoJ is an open-source imageJ pluggin (https://www.project-exoj.com)[8] based on the "à trous" wavelet transform (see Yuan *et al.*[1]) followed by spot tracking using the nearest-neighbour method. Fusion events are predicted according to a peak of fluorescence followed by an exponential decay. We used version 1.09 (2022-09-28) of ExoJ for our analysis. After several trials on out dataset, we determined values of the free parameters that produce satisfying results.

*Vesicle detection step*: the minimal and maximal vesicle radii were set to 2 and 5 pixels, respectively (pixel size is 160 nm). We did not apply photobleaching correction, because it created artifacts as mentioned by the authors.

*Tracking step*: the spatial searching range was set to 2 pixels. The temporal searching depth (gap closing) was set to 1 frame and the minimal event size was set to 2 frames. These values are given by default and were appropriate to process our data.

*Event detection*: we set to 5 the minimal number of points for fitting procedure, 4 the number of expanding frames (pre and post peak). We did not specify limits for the number of frames for decay, the estimated radius limit nor the maximum displacement.

The remaining parameters (wavelet scale σ, detection threshold dF/σ, minimal $R^2$ for decay and minimal $R^2$ for estimated radius) were more difficult to determine. Screening was limited by the fact that ExoJ is not a fully automatized algorithm. We run the analysis of the entire training dataset (60 movies) for 4 wavelet scales $\sigma \in \{2,3,5,10\}$. We applied a detection threshold dF/σ of 2, a minimal $R^2$ for decay of 0.7 (except for wavelet scale of 2 where we used 0.85 to reduce the number of movies where analyse could not be completed), and a minimal $R^2$ for radius of 0.6. Results table associates each event to dF/σ and two $R^2$, therefore higher thresholds could be tested *a posteriori* contrarily to wavelet scale that had to be set *a priori* (see **S1F**). We emphasize on the fact that a substantial part of the movies could not be analyzed (**S1E**); the event identification step never completed despite overnight computation. We tried different computers, different versions of ImageJ and to cut the movies without success. Most of the movies, which could not be analyzed, contained a vesicle detection step that was severely wrong, showing a high number of FPs that were even detected at high wavelet scales.

***Exocytosis detection with ADAE GUI***

Urbina *et al.* proposed an exocytosis detection method based on the h-dome transform, tracking with Kalman filtering and fitting of fluorescence intensity by an exponential decay[9]. Authors adapted the method in a new and open-source version of the MATLAB code called ADAE GUI[10] and we used it in our experimental study. The method detects exocytosis events using Difference of Gaussians (DoF). This algorithm, often applied for spot detection, consists in subtracting the result of the image filtered with a Gaussian filter of parameter $\sigma_1$ from the result of the image filtered with a Gaussian filter of parameter $\sigma_2>\sigma_1$. The resulting image presents sharp maxima at the spot locations. The first step of the algorithm consists in creating the mask of the cell. We skipped this step and put a unique cell mask for the whole movie as input, generated by thresholding the intensity (ImageJ plugin) of the first frame, because cell movements are minor in our 5-7min movies. This mask typically avoided false detection on the cell borders due to small movements of the cell. Next, a background subtraction (constant value defined as the average intensity outside the cell) was applied on the whole movie. Each frame was then analyzed separately, and two pre-processings were applied: an adjustment of the intensity histogram to match the histogram of the first frame (to correct potential bleaching) and a subtraction of the median projection of the five previous frames (to highlight exocytosis events and remove stationary objects and noise). DoG at several scales (9 scales in our case) was computed, with an initial $\sigma$ of 1 pixel ($\sigma$ is multiplied each time by 1.25). The selected image was the median projection of the 9 images. Applying this workflow to all frames, we obtained a movie with highlighted nonstationary spots. The maximum value in each frame was computed and the median of this set of values was defined as the threshold. If a spot had at least one pixel above this median value, it would be considered as an exocytosis event. Finally, a tracking algorithm (based on Kalman filter) was applied to avoid counting several times the same exocytosis event, if present in several successive frames. The exocytosis events were given as a CSV file containing the positions (x,y,t) of particles in the movie.

***Computation of performance scores***

The performances of the 3 methods (ExoDeepFinder, ADAE GUI and ExoJ) were evaluated in the same way through the comparison of 3D coordinates from predicted events with the ground truth. For each movie, a cell mask was generated (ImageJ intensity based thresholding on the first frame of the movie) and events outside the masks were removed before evaluation. Moreover, for the comparison of the 3 methods (**Figure 1**) (but not for robustness comparison) (**Figure 2**), events in the 25 first and last frames were removed to avoid any bias due to border effects. User can easily apply the same exclusions in real data. A predicted event was classified as a TP if the spatial distance $\Delta x$ to the nearest ground truth event is $\Delta x \leq 4.5$ pixels and the temporal distance $\Delta t$ is $-2 \leq \Delta t \leq 5$ frames (positive sign indicates predicted event after ground truth peak). These distances $\Delta x$ and $\Delta t$ for TP events are reported in **Figure S2B-C**. In the rare cases when several ground truth events matched the predicted event, the nearest spatial neighbor was chosen. All predicted events, which were not classified as TP, were classified as FP and the number of FN was computed as $\#FN = N - \#TP$ with $N$ the number of events in the ground truth list. The F1-score, Precision and Recall were computed as follows:

$$F1 - score = \frac{2 \times \#TP}{2 \times \#TP + \#FP + \#FN}$$

$$Precision = \frac{\#TP}{\#TP + \#FP}$$

$$Recall = \frac{\#TP}{\#TP + \#FN}$$

If $\#TP = 0$, $\#FP = 0$ and $\#FN = 0$ (once case in the Bafilomycin A1 dataset), these scores could not be computed.

##### *Lifetime and $F/F_0$ quantification*

Average fluorescence intensities were measured in a window of 7x7 pixels (1.12x1.12µm) centered on the exocytosis event during 16 frames (~5s) starting from peak intensity. The fluorescence intensity was normalized by the peak intensity. The intensity profile of all events in a cell was averaged. This averaged intensity decay was fitted with a single exponential function:

$$I(t) = Ae^{-t/t_{1/2}} + B$$

With $t_{1/2}$ the half-life (reported in **Table S3**). The fitting was performed with R using the minpack.lm function nlsLM().

The SBR ratio $F/F_0$ (reported in **Table S2**) was computed as the ratio between the mean intensity in a 7x7 pixels (1.12x1.12µm) window centered on exocytosis spot at the peak of fluorescence and 5 frames before. Contrarily to lifetime, a $F/F_0$ SBR was computed for each event and not averaged per cell.

##### *Venn diagram*

The construction of Venn diagrams (**Figure S2A**) started with the classifications of all events as TP, FP and FN for ExoDeepFinder, ExoJ and ADAE GUI. To be in intersection area, events associated to different detection methods were separated with $\Delta x \leq 4.5$ and $|\Delta t| \leq 7$ frames.

#### ExoDeepFinder

##### *Supervised deep learning for object detection in biological microscopy*

DeepFinder is an object detection method for 3D volumes that has originally been developed for detecting macromolecules in cryo-ET[11]. It is based on image segmentation using a 3D convolutional neural network, and is trained in a supervised manner. The network architecture is a U-net[12] and is described in[11]. After segmenting the objects of interest, the coordinates of the object centroid (*i.e.* center of mass) are retrieved by applying spatial clustering (mean-shift) to the segmentation map. Special features of ExoDeepFinder include an efficient implementation for handling large 3D datasets and strategies for dealing with rare classes. Furthermore, for training purposes, the user only needs to provide annotations in the form of a position list (*i.e.* coordinates of the objects centroid). ExoDeepFinder includes methods for converting the position list into an approximate segmentation map (to be used as training targets). The software is implemented in Python, is based on the tensorflow package, and is available at: https://github.com/deep-finder/tirfm-deepfinder.

##### *Custom optimization hyper-parameters*

To adapt DeepFinder (designed for cryo-ET) to TIRF microscopy, the distinctive characteristics of these image modalities needed to be considered. Cryo-ET data is a volume with 3 spatial dimensions, whereas TIRF data is a volume with 2 spatial dimensions and a temporal dimension (*i.e.* an image sequence). Therefore, we limited data augmentations like random rotations and mirroring to the (*x*,*y*) plane. Furthermore, from the model architecture perspective, we only applied maxpooling in the (*x,y*) dimensions. Since an exocytosis event typically lasts only 2 to 3 frames, it was essential to maintain a good temporal resolution. Finally, in terms of normalization, instead of using z-standardization (0-mean, 1-std) as used for cryo-ET, here we chose quantile normalization (so that the 1st percentile is aligned to value 0, and the 99th to value 1), because we noticed performance improvement for some image sequences.

***Generating segmentation maps from position lists:***

To train ExoDeepFinder, only the objects position (centroid coordinate) needed to be annotated. Segmentation maps for training were then generated by placing an (*x,y,t*) shape model at annotated positions. The luminescence of an exocytosis is isotropic in the (*x,y*) plane (*i.e.,* a disc) and has an exponential decay in *t*. Therefore our (*x,y,t*) shape model was a tube of 3 frames with an exponential decaying radius (starting at R=4 pixels, then R=2 and ending at R=1 pixel).

***Handling rare events***

The class imbalance is severe in our target case study: most image sequences have less than 0.1% of pixels that belong to the exocytosis class. To deal with this problem, ExoDeepFinder samples 3D patches centered on annotated positions. The proportion of exocytosis event pixels in the patch is then more favorable for achieving a successful training. In this respect, when choosing the patch size, there is a trade-off between class imbalance and detection performance. On the one hand, a small patch size results in a higher proportion of the rare class, because the patch is wrapped tighter around the object of interest. The downside is that less context is included in the model decision making. As a result, the model has a high false positive rate, as the missing context of the object is an important cue. On the other hand, a large patch size mitigates this problem, as with more context the model disposes of more information, hence a lower false positive rate. Yet, the proportion of the rare class within the patch is too low, which causes the training to fail (the cost function does not decrease and the model always predicts the over-represented class, *i.e.* the negative class). We solve this trade-off by modulating the patch size during training, from small (83 pixels) to large (483 pixels). Starting with a small patch size ensures a good proportion of exocytosis event pixels (12.3% for 83 patches), allowing to obtain initial model weight values that will ensure the success of the subsequent training. We then increase the patch size by a factor 2 every 10k training iterations (see parameters in **table S4**), so that the model gradually incorporates more image context to its decision making, which effectively decreases the false positive rate (that is initially high for small patch sizes).

***Enhance detection performance by providing counter-examples***

Furthermore, we noticed that ExoDeepFinder tends to confuse exocytosis events (*i.e.,* blinking spots) with docked vesicles (*i.e.,* spots with no exponential decay, whose luminescence remains constant through frames). We therefore decided to include counter-examples to the training procedure. We chose to generate these counter-examples by using the unsupervised spot detector ATLAS[13], which is avalaible at: https://gitlab.inria.fr/serpico/atlas. ATLAS software enables to detect spots in 2D fluorescence images. The spot size is automatically selected and the detection threshold adapts to the local image dynamics. ATLAS relies on the Laplacian of Gaussian (LoG) filter, which both reduces noise and enhances spots. A multiscale representation of the image is built to automatically select the optimal LoG variance. Local statistics of the LoG image are estimated in a Gaussian window, and the detection threshold is pointwise inferred from a probability of false alarm (PFA). The user only has to specify i) standard deviation of the Gaussian window and ii) PFA value. The Gaussian window must be about the size of the background structures; increasing the PFA increases the number of detections. In all our experiments, we set PFA to 0.001 and the standard deviation of the Gaussian window to 21 pixels. All connected components lower than 3 pixels are discarded. Then, we merge the Atlas detections with the exocytosis event annotations into a multi-class segmentation map (label 0 is background, label 1 is exocytosis event, label 2 is docked vesicle). As the Atlas detections were not supervised by an expert, they contain some FPs and FNs. The quality of these detections was nevertheless sufficient for our purpose (*i.e.,* to provide counter examples). In practice, it enhanced the detection performance for exocytosis events, and it also helped to mitigate the imbalanced class problem, as shown in[11].

***Additional implementation details and computing times***

ExoDeepFinder has been computationally trained with the Tversky loss[14] and ADAM algorithm, chosen for its good convergence rate, by setting the learning rate to 0.0001, the exponential decay rate to 0.9 for the first moment estimate and to 0.999 for the second moment estimate. No regularization (e.g., L2 regularizer or 'drop out') was used for processing the datasets.
The inference stage of ExoDeeFinder is fast as it takes only 33.79 seconds using a Nvidia A100 GPU to process a movie of 1001 images of 261 x 149 pixels. The training from hybrid annotations required 17 hours and 30 minutes on a Nvidia A100 GPU.

### Datasets

The training/inference dataset is composed of 120 movies of single RPE1 VAMP7-pHluorin transfected cells acquired in TIRFM. Part of this dataset is already published in Lachuer et al.[7,5]. All these cells are in control conditions (no drug treatment). However this training dataset includes variation in the seeding conditions. While most of the cells are seeded on fibronectin-coated glass-bottom dish (62 movies), some of them are seeded on PLL (23 movies) or micropatterns (ring or rectangles shaped) (17 and 18 movies respectively). Moreover some of the conditions are co-transfected, either with Paxillin-mCh (19 movies) or mCh-Rab6A (5 movies). We include this diversity in the dataset to increase the robustness of ExoDeepFinder. The histamine and Bafilomycine A1 datasets come also from Lachuer et al.[5]. The HeLa and CD63 datasets were not published elsewhere. While HeLa cells were seeded on fibronectin coated glass, CD63-pHluorin transfected RPE1 cells were seeded on ring-shaped micropattern.
The whole dataset is publically available at https://doi.org/10.5281/zenodo.11204932.

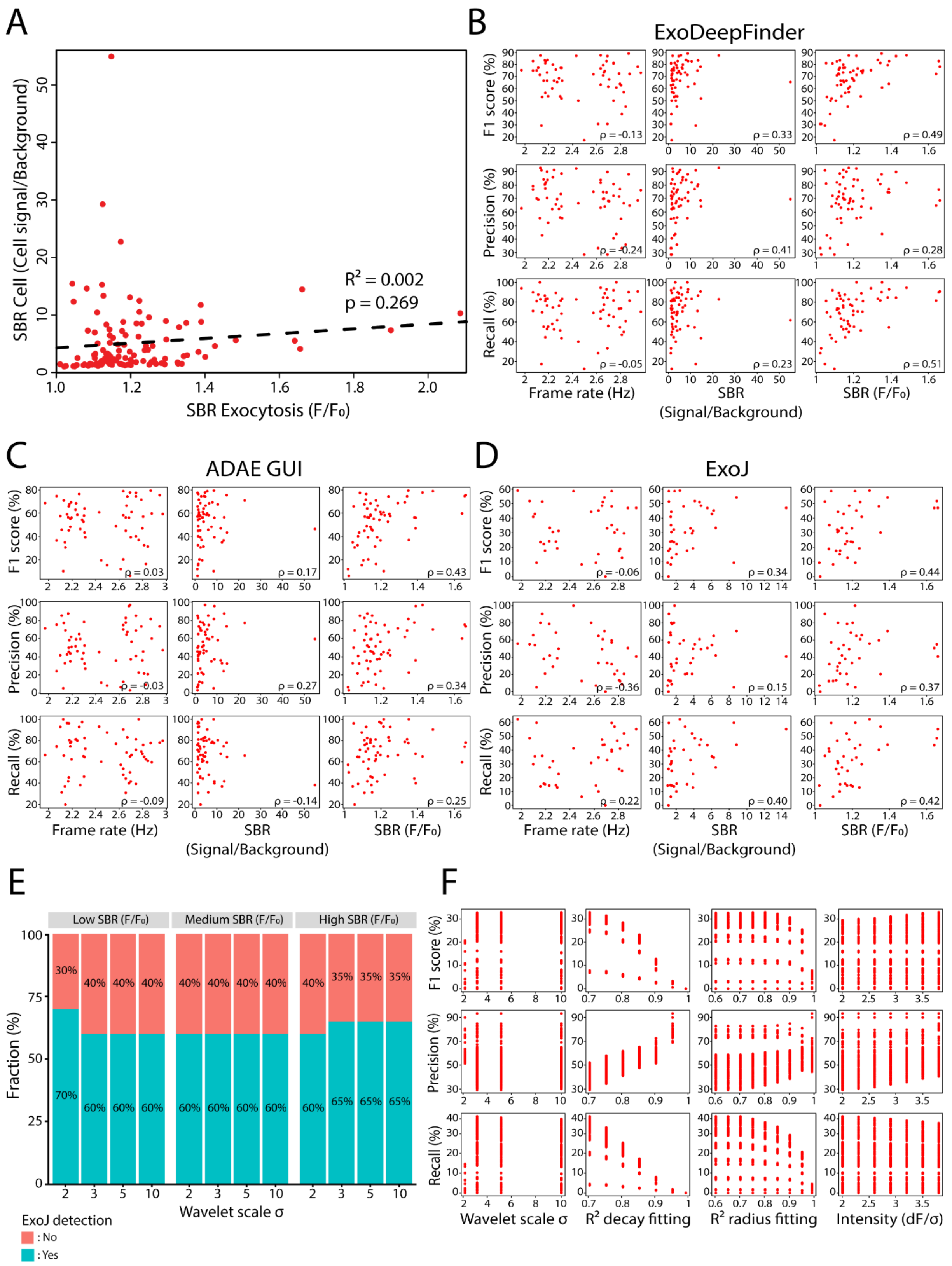


**Figure S1**. **A**. Correlation between the two types of SBR (cell signal/background and F/F$_0$) in the training dataset (60 movies). Significance of the correlation has been evaluated through a t-test and the $R^2$ is indicated. **B**. Correlation of ExoDeepFinder performances with frame rate and SBR (cell signal/background and F/F$_0$). Spearman correlation coefficients are indicated for each plot. **C**. Correlation of ADAE GUI performances with frame rate and SBR (signal/background and F/F$_0$).

Spearman correlation coefficients are indicated for each plot. **D**. Correlation of ExoJ performances with frame rate and SBR (cell signal/background and $F/F_0$). Spearman correlation coefficients are indicated for each plot. **E**. Fraction of inference dataset for which ExoJ analysis is possible as a function of the different wavelet scales used. **F**. ExoJ performances on the training dataset as a function of its different parameters.

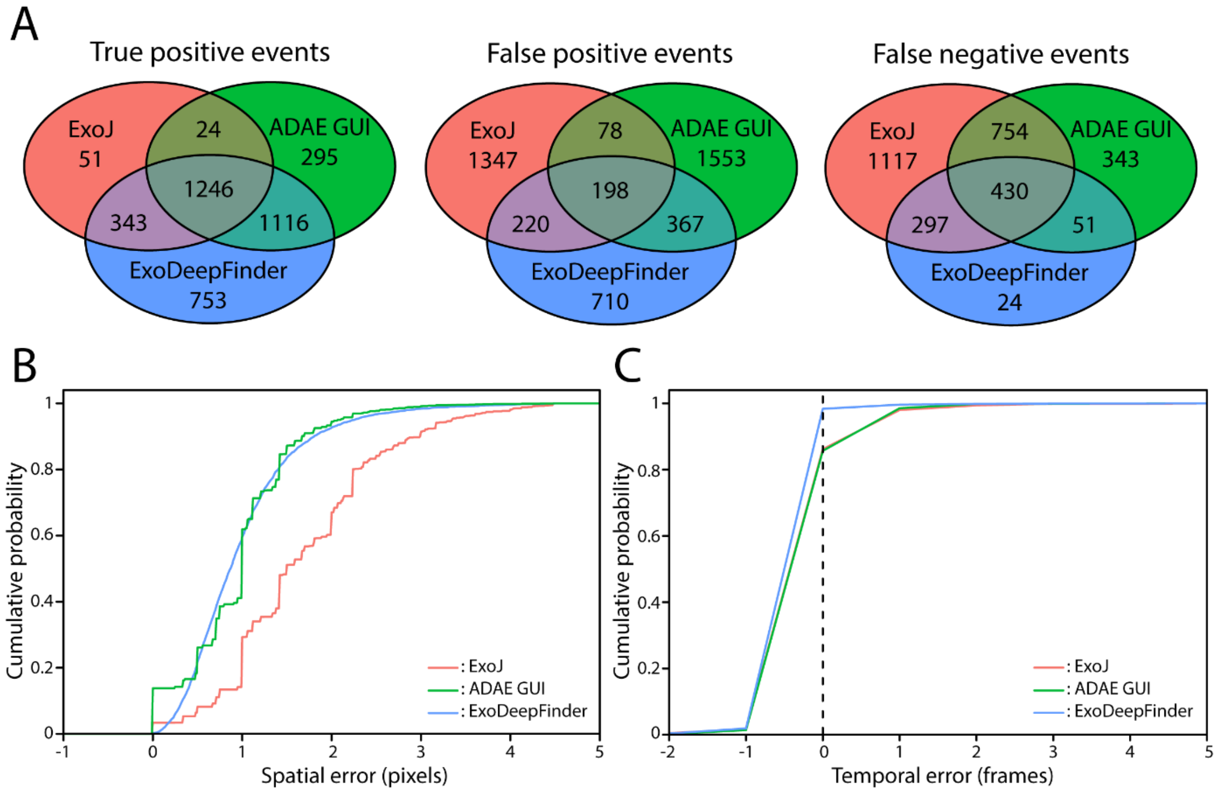


**Figure S2**. **A**. Venn diagram of TP, FP and FN events. Numbers indicated are the absolute counts in each category. **B**. Cumulative distribution of spatial localization error for TP events detected by ExoJ, ADAE GUI and ExoDeepFinder. Note that the different smoothness of the curves is due to the different strategies of sub-pixel localization. **C**. Cumulative distribution of temporal localization error for all true positive events detected by ExoJ, ADAE GUI and ExoDeepFinder. In B and C, number of events for each curve are $n_{ExoJ}$ = 1664, $n_{ADAE\ GUI}$ = 2681 and $n_{ExoDeepFinder}$ = 3458. In B-C, ExoJ has less number of movies analyzed (hence less total number of events), because the analysis could not be performed for 30-40% of the data (**see method**). In B-C, ExoDeepFinder was trained on the total 60 movies of the training dataset (model all).

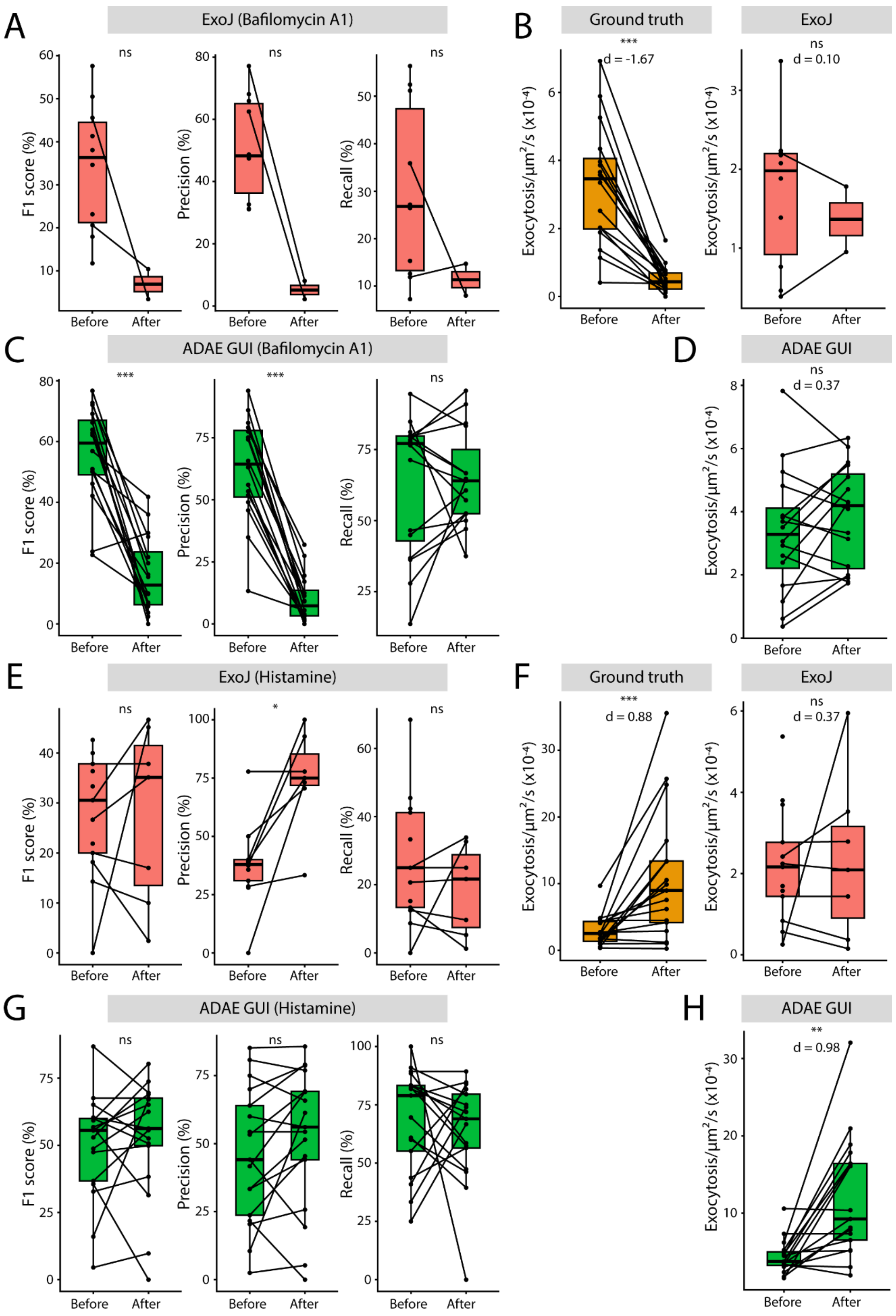


**Figure S3**. **A**. ExoJ performances before and after Bafilomycin A1 (100nM, 60 min) treatment. **B**. Exocytosis rate before and after Bafilomycin A1 (100nM, 60 min) treatment measured by manual detection (ground truth) and compared to ExoJ detection. Note that in A-B, unpaired points are due to movies for which ExoJ analysis was not possible. In A-B, n=16 cells from three independent experiments for ground truth but only 2 cells for which analysis was possible before and after treatment. **C**. ADAE GUI performances before and after Bafilomycin A1 (100nM, 60 min) treatment. **D**. Exocytosis rate before and after Bafilomycin A1 (100 nM, 60 min) treatment measured by manual detection (ground truth) and compared to ADAE GUI detection. In C-D, n=16 cells from three independent experiments. **E**. ExoJ performances before and after histamine (100µM, cells immediately imaged) treatment. **F**. Exocytosis rate before and after histamine (100µM, cells immediately imaged) treatment measured by manual detection (ground truth) and compared to ExoJ detection. Note that in E-F, unpaired points are due to movies for which ExoJ analysis was not possible. In E-F, n=17 cells from three independent experiments for ground truth but only 6 cells for which analysis was possible before and after treatment. **G**. ADAE GUI performances before and after histamine (100µM, cells immediately imaged) treatment. **H**. Exocytosis rate before and after histamine (100µM, cells immediately imaged) treatment measured by manual detection (ground truth) and compared to ADAE GUI detection. In G-H, n=17 cells from three independent experiments. In B,D, F and H, significance has been evaluated with paired Wilcoxon test, ns $p>0.05$, *$p<0.05$, **$p<0.01$ and ***$p<0.001$. In B, D, F and H, effect sizes are measured with the Cohen's *d* for paired samples.

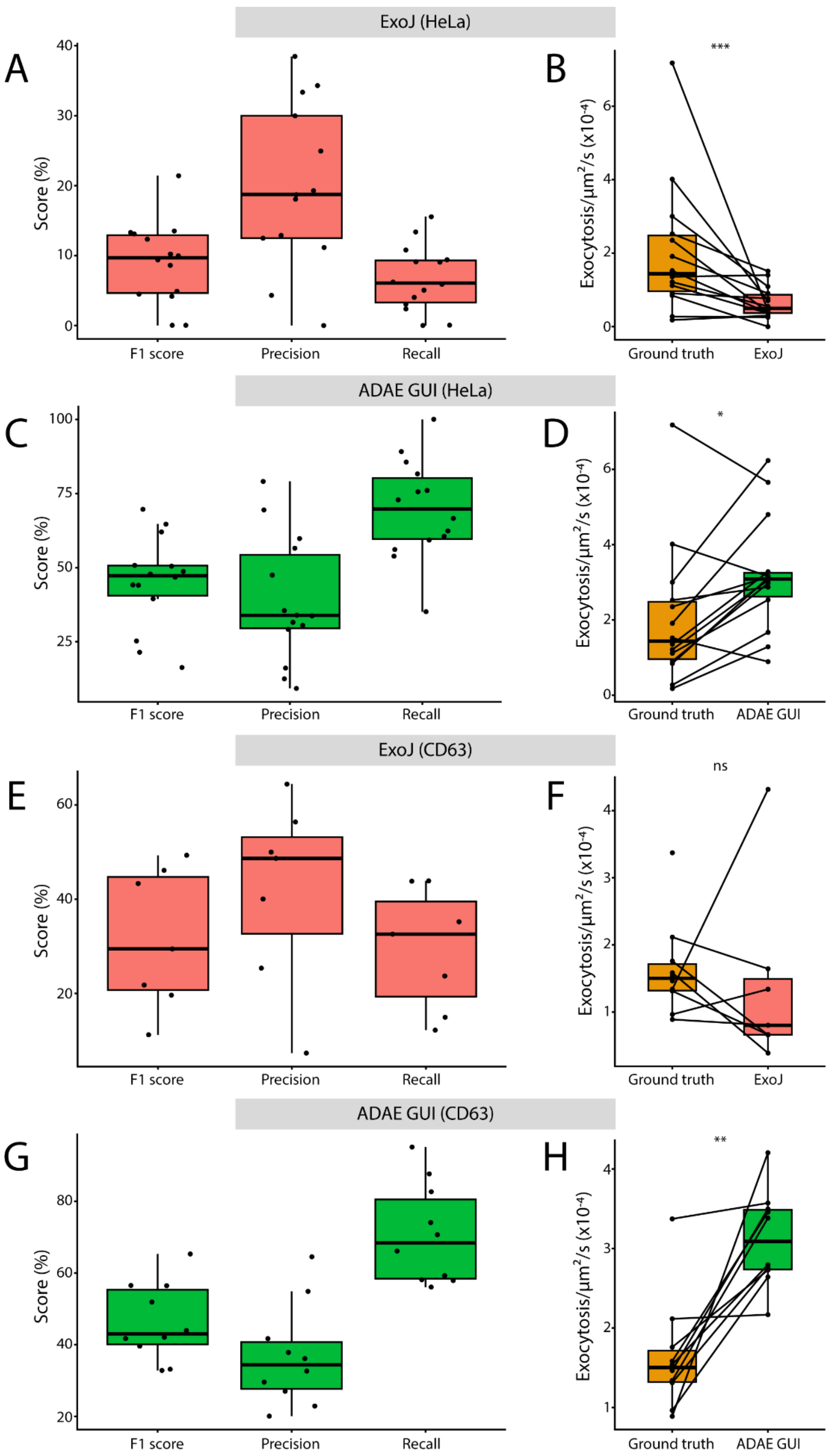


**Figure S4**. **A**. ExoJ performances in VAMP7-pHluorin transfected HeLa cells. **B**. Comparison of the exocytosis rate measured by manual detection (ground truth) and compared to ExoJ detection. In A and B, 14 cells analyzed from a single experiment (all movies could be analyzed with ExoJ). **C**. ADAE GUI performances in VAMP7-pHluorin transfected HeLa cells. **D**. Comparison of the exocytosis rate measured by manual detection (ground truth) and compared to ADAE detection. In C and D, 14 cells analyzed from a single experiment. **E**. ExoJ performances in CD63-pHluorin transfected RPE1 cells. **F**. Comparison of the exocytosis rate measured by manual detection (ground truth) and compared to ExoJ detection. In E-F, n=10 cells from a single experiment for ground truth but only 6 cells for which ExoJ analysis was possible. **G**. ADAE GUI performances in CD63-pHluorin transfected RPE1 cells. **H**. Comparison of the exocytosis rate measured by manual detection (ground truth) and compared to ADAE GUI detection. In G and H, 10 cells analyzed from a single experiment. In B, D, F and H, significance has been evaluated with paired Wilcoxon test, ns $p>0.05$, *$p<0.05$, **$p<0.01$ and ***$p<0.001$.

### Movies legends

**Movie S1**: Representative movie of a VAMP7-pHluorin transfected RPE1 cell in TIRFM. White squares highlight some of the exocytosis events characterized by a sudden apparition of a bright spot and then the 2D diffusion of the signal. Time code is in minute:second format.

**Movie S2:** Representative movie of a VAMP7-pHluorin transfected RPE1 cell in TIRFM showing the detection of the exocytosis events for the 3 described methods, compared with the ground truth. Each event appears 3 frames before and remains 3 frames after. Performances were compared from frame 20 (00:19) to frame 981 (16:20).

### Tables

| | | Subset A | Subset B | Subset C | Subset D | Subset E | All |
|---|---|---|---|---|---|---|---|
| **# of exocytosis events** | **Total** | 630 | 1320 | 2870 | 5732 | 8698 | 11898 |
| | Low SBR | 227 (36.03%) | 515 (39.02%) | 1257 (43.80%) | 2522 (44.00%) | 3872 (44.52%) | 5458 (45.87%) |
| | Medium SBR | 158 (25.08%) | 385 (29.17%) | 773 (26.93%) | 1530 (26.69%) | 2290 (26.33%) | 3064 (25.75%) |
| | High SBR | 218 (34.60%) | 420 (31.82%) | 840 (29.27%) | 1680 (29.31%) | 2536 (29.16%) | 3376 (28.37%) |
| **# of movies** | **Total** | 8 | 10 | 20 | 38 | 53 | 60 |
| | Low SBR | 4 (50.00%) | 2 (20.00%) | 7 (35.00%) | 13 (34.21%) | 18 (33.96%) | 20 (33.33%) |
| | Medium SBR | 2 (25.00%) | 4 (40.00%) | 7 (35.00%) | 13 (34.21%) | 17 (32.08%) | 20 (33.33%) |
| | High SBR | 2 (25.00%) | 4 (40.00%) | 6 (30.00%) | 12 (31.58%) | 18 (33.96%) | 20 (33.33%) |

**Table S1**. Description of the different ExoDeepFinder training datasets of in terms of absolute number of exocytosis events and number of movies. For each training dataset, the numbers of exocytosis events and movies coming from the 3 originally defined training datasets (low, medium and high SNR) are indicated. Fraction of the total numbers are indicated in brackets.

| | ExoJ | ADAE GUI | ExoDeepFinder |
|---|---|---|---|
| **TP** | 1.38 ± 0.35 | 1.33 ± 0.28 | 1.29 ± 0.26 |
| **FP** | 1.11 ± 0.15 | 1.16 ± 0.13 | 1.17 ± 0.14 |
| **FN** | 1.22 ± 0.19 | 1.16 ± 0.15 | 1.13 ± 0.11 |

**Table S2**. SBR ($F/F_0$) for each class of event (TP, FP and FN) for the different detections methods. Values are means (over all events) ± SD.

| | ExoJ (s) | ADAE GUI (s) | ExoDeepFinder (s) |
|---|---|---|---|
| **TP** | 2.00 ± 1.08 | 2.03 ± 1.12 | 2.05 ± 1.12 |
| **FP** | 1.17 ± 1.67 | 2.13 ± 2.46 | 1.98 ± 1.42 |
| **FN** | 1.71 ± 1.29 | 1.54 ± 1.77 | 1.35 ± 0.93 |

**Table S3**. Lifetime of exponential decay of exocytosis event for each class of event (TP, FP and FN) for the different detections methods. Values are means (over all cells) ± SD. Note that contrarily to SBR in **Table S2**, a single lifetime is computed per movie. Several events need to be merged to obtain a robust estimation of the decay contrarily to SBR that can be easily evaluated for each single event.

| | Training iterations | Patch size (in pixels) | Maximal random shift (in pixels) | Batch size |
|---|---|---|---|---|
| **Round 1** | 0-10k | 8 | 4 | 256 |
| **Round 2** | 10k-20k | 16 | 8 | 128 |
| **Round 3** | 20k-30k | 32 | 16 | 32 |
| **Round 4** | 30k – Until convergence | 48 | 32 | 10 |

**Table S4**. Training schedule. For its round, the parameters has been changed. The patch size determines the size of the local context. A random shift is applied to the patch sampling as a data augmentation technique. Hence, objects of interest are not always located at the center of the patch[11]. The batch size is chosen according to the available GPU capacity.
